# Forecasting climate-driven distributional changes in the threatened Caribbean marine species *Aliger gigas* (Queen conch)

**DOI:** 10.64898/2026.04.29.721193

**Authors:** Daniel Rojas-Ariza, Claudia Nuñez-Penichet, Zenia Ruiz-Utrilla

**Affiliations:** Department of Ecology and Evolutionary Biology, University of Kansas, United States; Biodiversity Institute, University of Kansas, United States; Fish and Wildlife Conservation Department, Virginia Tech, United States

**Keywords:** Climate change, distributional ecology, ecological niche modeling, marine fisheries, threatened species

## Abstract

The queen conch (*Aliger gigas*) is a key native species of the Caribbean Sea and a primary source of income for thousands of fishers. Historically, it has been a highly valuable resource for the fishing sectors of countries such as the Bahamas, Turks and Caicos, Honduras, and Nicaragua. However, due to its high economic value, the species has been extensively overfished across the region. Overfishing, combined with limited larval dispersal, low recruitment, and poor population connectivity, has led to a drastic decline in population numbers of the species, resulting in its current classification as Threatened. Despite this status, likely impacts of climate change on its populations remain poorly understood, posing significant challenges to conservation efforts. To address this gap, we integrated occurrence records, climate data, and satellite-derived marine habitat data to develop ecological niche models estimating the current and future distribution of the queen conch under different climate change scenarios. We found substantial losses of suitable areas for queen conch along the northern Atlantic coast of South America and Central America, part of the Greater Antilles and the Lesser Antilles. The entire Caribbean region is projected to lose suitability entirely within 20-30 years under the moderate and most extreme climate scenarios. Conversely, our models estimate some suitable areas to persist or expand along the southeastern coast of the United States at least until sometime between 2040 and 2060. Overall, our results suggest a northward shift in the range of this species, with the magnitude of this shift closely tied to the severity of climate change impacts. This work aims to build upon and enhance existing knowledge about survival of queen conch populations in the Caribbean over time. Anticipating future habitat availability will be key to protecting this economically and ecologically important species.

## 1 Introduction

The queen conch, *Aliger gigas* (Linné and Salvius 1758), is a charismatic species of the Western Atlantic that has played an important role in the history of the Caribbean region and human communities of the West Indies over the past several centuries (Brownell and Stevely 1981; Thelie 2001; Antczak and Antczak 2005; Antczak et al. 2008 and references therein) and represents an important food and shell source for more than 30 nations in the region (Stoner et al. 2023). This species is distributed throughout the Gulf of Mexico, the Caribbean Sea, Bermuda, and the coast of South America (Clench and Abbott 1941; Thelie 2001), north of the mouth of the Orinoco River in Venezuela (Tewfik 1996).

Historically, this species has been important in the fishing sector of the Caribbean, and in supporting livelihoods of various communities in the West Indies. It was classified as the second most valuable Caribbean fishery resource (after spiny lobster, Panulirus argus) in 1981 by Brownell and Stevely (1981), and it has served as the primary source of income for ∼20,000 fishers in the region (Prada et al. 2017). The most economically important product derived from this species is its meat (Horn et al. 2022), with an estimated value of US$60M in 2003 (Thelie 2003) and ∼US$74M in 2017 (Prada et al. 2017). Most of the catch is landed in the Bahamas, Turks and Caicos, Honduras, Nicaragua, Belize, and Jamaica (Clench and Abbott 1941; Horn et al. 2022). Turks and Caicos, Bahamas, and Honduras have been the leading producers of queen conch since the 1950s, and Jamaica and Nicaragua have emerged as major producers only since the 1980s and 2000s, respectively (Sea Around Us Program, SAU, Pauly et al. 2020). In 2013, Nicaragua had the highest production (US$9.0M), followed by Belize (US$5.5M), Turks and Caicos Islands (US$3.8M), and the Bahamas (US$3.2M; Prada et al. 2017).

This species has been found in a variety of habitats, depending on its different life stages (Stoner 2003; Horn et al. 2022). As juveniles, queen conchs are most commonly associated with seagrass ecosystems (Alcolado 1976; Hesse 1979; Stoner 2003; Stoner et al. 1996; Stoner et al. 1994; Weil and Laughlin 1984). Metamorphosis of queen conch veligers is triggered by trophic signals in seagrass habitats (Davis et al. 2005; Davis and Stoner 1994), which may explain why these ecosystems are key nursery habitats for the species (Stoner et al. 1994; Stoner 2003). Nevertheless, juveniles have also been reported in other habitats, including coral rubble and sand flats (Glazer and Berg 1994; Randall 1964; Torres Rosado 1987). As adults, they are found in a broader range of habitats, including sandy bottoms and algal or coral rubble (Acosta 2001; Stoner and Davis 2010), and at depths less than 75m, but most commonly <30m (McCarthy 2007).

Although several studies have documented and predicted loss and degradation of critical seagrass habitats (e.g., Duarte 2002; Orth et al. 2006; Vallés and Oxenford 2012; van Tussenbroek et al. 2014; Waycott et al. 2009), no conclusive evidence shows that habitat modification, contraction, or disappearance within the range of queen conch is significantly contributing to its extinction risk (Horn et al. 2022). The broad range of habitats occupied by queen conch at different life stages, along with their capacity to use highly variable marginal habitats (Dujon et al. 2019; Stieglitz et al. 2020), may contribute to the species’ resilience against local extinction. Overfishing, on the other hand, has been identified as a key cause of sustained declines in queen conch abundance and the collapse of several populations across its range in recent decades (Horn et al. 2022; Prada et al. 2017; Stoner et al. 2018; WECAF 1980). Low population densities, lack of population connectivity in some areas, limited larval dispersal, and low recruitment rates have led the National Marine Fisheries Service (NMFS) of the United States to classify it as Threatened under the Endangered Species Act (ESA, NMFS, 2024). The species is considered to be at a moderate risk of extinction, meaning that it “is on a trajectory that puts it at high risk of extinction over the next 30 years” (Horn et al. 2022, p. 103).

Another factor of potential threat for queen conch may be the future changes in sea temperatures, as these have been reported to influence the species’ reproduction cycles, egg and larval development, larval survival rates (e.g., Davis 2000; Aldana Aranda and Manzano 2017; Chavez Villegas et al. 2017) and, to a lesser extent, shell biomineralization processes (Aldana Aranda and Manzano 2017). By 2100, mean sea surface temperatures (SSTs) are projected to increase between 1.1 and 6.4°C globally (Intergovernmental Panel on Climate Change 2007), and between 1.9 and 3.0°C in the Caribbean (Bustos Usta and Torres Parra 2021). Identifying present and future environmental suitability across the Caribbean for queen conch, in view of these significant environmental changes, is critical to anticipate emerging threats and guide conservation planning. Given the current conservation status of queen conch and the dynamic environmental context, it is crucial to investigate and predict, to the greatest extent possible, future distribution patterns of this species under likely scenarios of climate change. In this study, we estimated likely changes in the distributional potential of queen conch in the coming decades across different climate change scenarios and evaluated the environmental suitability of different areas of the Caribbean relative to the presence of seagrass habitat and marine protected areas (MPAs).

## 2 Materials and Methods

### 2.1 Occurrence records and environmental variables

We downloaded occurrence records for queen conch from the Global Biodiversity Information Facility database (GBIF 2024) and the Ocean Biodiversity Information System (OBIS) using the R packages rgbif (v. 3.8.0, Chamberlain 2024) and robis (v. 2.11.3, Provoost and Bosch 2022), respectively. For the GBIF dataset, we retained only records that had geographic coordinates (longitude and latitude) recorded to at least two decimal places, associated with specimens preserved in collecting or research institutions (e.g., museums and research centers). For the OBIS data, we also retained observational records, provided they were associated with an institution. We filtered the data by removing (1) dubious records located outside of the known distribution of the species in the Caribbean region (Thelie, 2001), (2) duplicate records (i.e., identical latitude and longitude), and (3) records lacking year information (Cobos et al. 2018). To further reduce uncertainty in our suitability estimates and avoid introducing potential erroneous signals, we also removed records located outside of the known depth range of queen conch (<75 m, McCarthy 2007), using a dataset from the General Bathymetric Chart of the Oceans (15-arc seconds spatial resolution, GEBCO Compilation Group 2025). This range is thought to be constrained by the photosynthetic zone of the sea, which limits the food sources available to the species (Robertson 1962). Additionally, to reduce spatial autocorrelation, we applied a distance-based thinning procedure, retaining only one record per environmental pixel (see details on environmental variables below), following Anderson (2012). These thinned records were then divided into two subsets: 70% of the records were used for model training, and the remaining 30% for model evaluation.

We considered six ocean surface environmental variables for ecological niche models: ocean temperature, salinity, seawater velocity and direction, primary productivity, and pH. These variables were obtained from the Bio-ORACLE database (v3.0, 0.05° spatial resolution, Assis et al. 2024) using the R package biooracler (v. 0.0.0.9000, Fernandez 2024). Each variable is represented by six decadal statistics: maximum, minimum, average, average of the yearly maxima, average of the yearly minima, and range, for a total of 36 environmental layers per decade. We averaged the values of each environmental variable over 20-year intervals between 2000 and 2080. The average values for 2000-2020 represented the present-day scenario, and were used for model calibration; while the 2020-2040, 2040-2060, and 2060-2080 represented future predicted conditions, and were used for model transfer. For the future projections, we considered three Shared Socioeconomic Pathways (SSP, Riahi et al. 2017; van Vuuren et al. 2017) to include different scenarios of carbon emission: low (SSP-126), moderate (SSP-370), and high (SSP-585). To reduce dimensionality and avoid multicollinearity, we performed a principal component analysis (PCA) on the 2000-2020 raw environmental variables using the kuenm2 R package (v. 0.1.1, Tridade et al. 2025). The first five PCs were selected for modeling by the algorithm, which together accounted for ∼79.4% of the variance of the original environmental data. We then applied those principal components equations to each combination of SSP scenario and 20-year period (e.g., SSP 126 during 2040–2060). PC layers were masked to the area of interest (see Figure 1) using a reference shapefile of the Atlantic Ocean from marineregions.org (Flanders Marine Institute, 2024).

**Figure 1.**
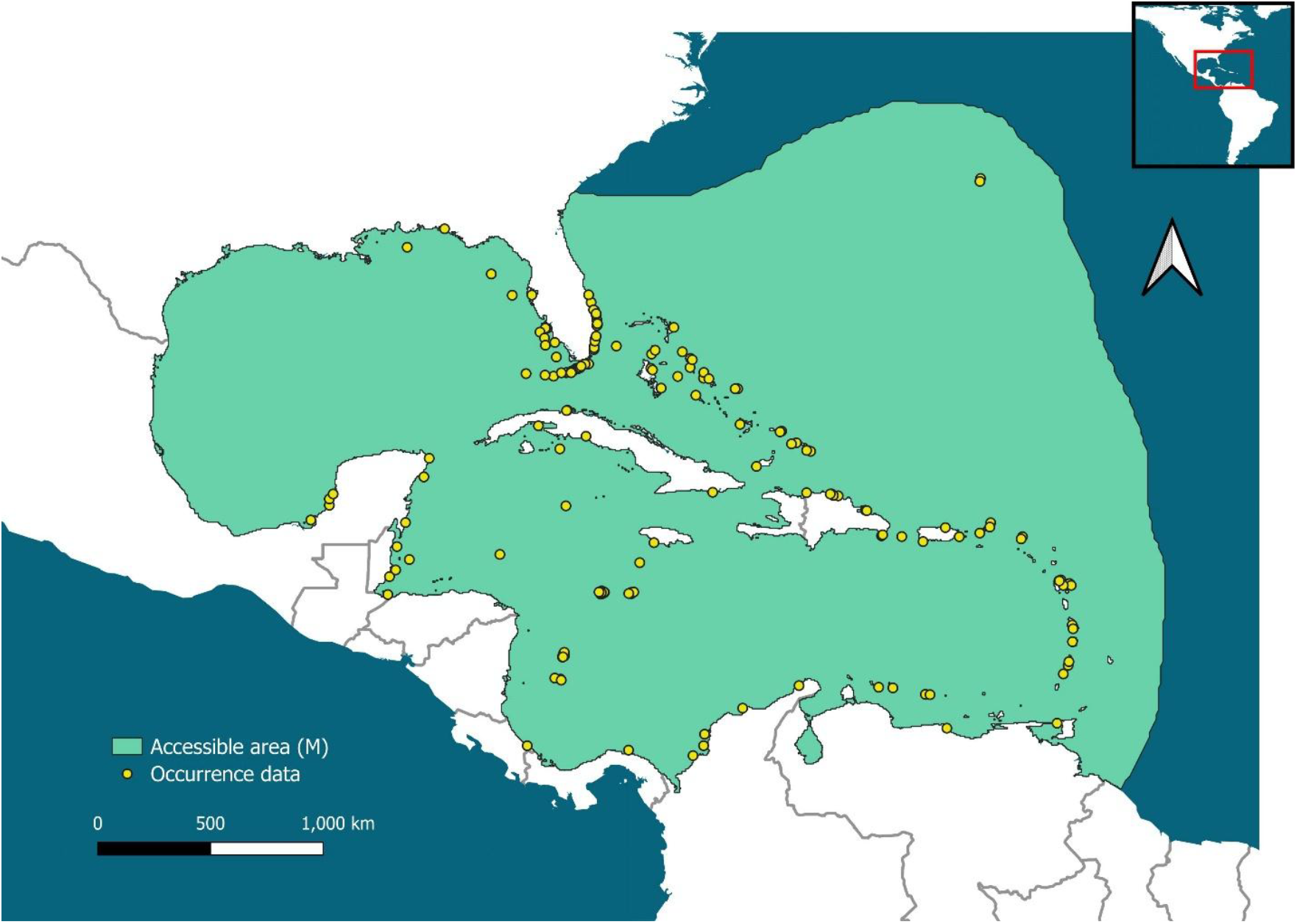
Occurrence records and hypothesized accessible area (**M**; calibration area) of the queen conch (*Aliger gigas*) used in ecological niche models.

### 2.2 Ecological Niche Modeling and Future Projections

Two algorithms were used to estimate ecological niches of the queen conch: Maxent (Phillips et al. 2017) and ellipsoids (Cobos et al. 2022). Employing both algorithms enables us to evaluate the consistency of our predictions with and without consideration of a hypothesized accessible area and environmental background. All data preparation and modeling processes were executed in R (R Core Team 2025).

We used Maxent (Phillips et al. 2017) to calibrate ecological niche models and generate candidate models. Together, the Caribbean Sea, the Gulf of Mexico, and the western part of the Sargasso Sea were considered as the accessible areas for ecological niche models developed with Maxent (M; Soberón and Peterson 2005, see Figure 1). To account for potential sampling bias in queen conch presence data, we generated a bias file for use during Maxent model calibration (see Trindade et al., 2026 for documentation on bias file implementation). We used GBIF occurrence records of marine gastropods from the Caribbean (GBIF 2026) as a surrogate for spatial sampling effort, under the assumption that all species in this target group are collected using the same survey methods as queen conch (Phillips et al. 2009; Anderson 2012). These occurrence records were subjected to the same data cleaning and spatial thinning procedures applied to the queen conch GBIF dataset described above. Point counts were rasterized and smoothed using a Gaussian kernel to produce a continuous raster of probability weights, representing the relative likelihood of sampling at any given location across the study area. This bias raster was incorporated into Maxent to weigh the selection of environmental background points proportionally to sampling effort, reducing the influence of spatially uneven data collection on model outputs (Phillips et al. 2009; Anderson 2012).

We produced 1170 candidate models from combinations of the following parameters: 5 feature classes (q, lq, lp, qp, lqp; l: linear, q: quadratic, and p: product), 9 regularization multipliers (0.10, 0.25, 0.50, 0.75, 1.00, 2.00, 3.00, 4.00, 5.00), and 26 combinations of two or more of the selected PCs of environmental variables. We used three metrics to assess model performance and model selection. First, we evaluated the statistical significance of predictions of the species’ distribution using partial ROC (Peterson et al. 2008) and assessed predictive performance via omission rates at a 5% training presence threshold (E = 5%; Anderson et al. 2003). We finally evaluated model complexity using the Akaike Information Criterion corrected for small samples (AICc, Warren and Seifert 2011). A common assumption in ecological niche theory is that responses of species to environmental gradients show unimodal response curves (Austin 2007; Hutchinson 1957). Although bimodal response curves can certainly be generated in model fitting (e.g., a quadratic response with upward concavity), they are inconsistent with expectations of ecological niche theory (Austin 2007, Qiao and Escobar 2025), and therefore models with bimodal responses were discarded. Model calibration and selection were done using the R package kuenm2.

Models retained after the selection process were projected to the full extent of the Caribbean Sea for the current environmental context and the future 20-year climatic SSPs scenarios. We generated 10 bootstrap replicates for each model under three different extrapolation types available in Maxent (NE: no extrapolation, EC: extrapolation with clamping, E: free extrapolation). Our analysis focuses on predictions under the free extrapolation scheme, as it produces models consistent with the assumption of unimodal-shaped species responses to environmental gradients (Hutchinson 1957; Austin 2007). If more than one model met the selection criteria, we generated a consensus by calculating the mean suitability values across those models for each scenario, using the kuenm2 package. The resulting suitability estimates were then binarized (i.e., converted to presence/absence) using model-specific thresholds calculated using the omission rate defined above. Binarizations were filtered to retain only areas shallower than 75m to refine habitat estimates to depths occupied by the species (corresponding to the typical depth range of queen conch Randall 1964; McCarthy 2007). This filtering was performed using a higher-resolution bathymetric dataset from the General Bathymetric Chart of the Oceans (GEBCO; 15-arc seconds spatial resolution; GEBCO Compilation Group, 2025).

We also generated ecological niche models based on ellipsoids using the ellipsenm package in R (v.0.3.4, Cobos et al. 2022). We calibrated and evaluated 26 ellipsoid models based on covariance matrices produced from the combinations of PCs representing the environmental variables to identify and select the models that yielded the lowest omission rates and lowest prevalence among all those analyzed (i.e., the proportion of geographic and environmental space predicted as suitable, see Cobos et al. 2022 for details on ellipse calculation method). To account for potential biases in the occurrence data, we generated five ellipsoid envelopes for each best-supported model based on bootstrapped subsamples of 75% of the data and a 95% confidence level for the pairwise confidence region of the ellipsoids. Suitability estimates were subsequently binarized using a 5% omission threshold (aligned with the confidence level) and were filtered to the range of depths for this species following the same methods described for Maxent models above.

### 2.3 Suitability estimates within seagrass habitats in the Caribbean

Seagrass beds are key habitats for juvenile queen conch (Stoner 2003), making it important to evaluate ecological niche model outputs with a focus on these critical environments. To this end, we identified how much of the estimates of current suitable areas (2000–2020) overlapped with the distribution of seagrass habitats in the region, using the known extent of benthic seagrass beds mapped by the Allen Coral Atlas for the Caribbean (Allen Coral Atlas, 2022). Because our model output raster had a coarser resolution (15-arc seconds) than the seagrass vector data, we considered a pixel of a suitability projection as overlapping if it contained at least one seagrass polygon. We then quantified the proportion of pixels containing seagrass beds that were predicted as suitable by our ecological niche models.

### 2.4 Suitability estimations within Marine Protected Areas (MPAs) in the Caribbean

To evaluate potential changes in environmental suitability relative to the current marine protected areas (MPAs) in the Caribbean, we estimated the proportion of currently abiotically suitable areas for queen conch covered by MPAs. We obtained shapefiles of these MPAs defined worldwide from the World Database on Protected Areas (WDPA, UNEP-WCMC, and IUCN 2024), and cropped them to the Caribbean region (Supplementary Figure S1). We estimated the proportion of suitable areas in our present-day distribution estimates covered by the current MPA configuration in the region. Additionally, these estimates were assessed in relation to the configuration of MPAs across four Caribbean areas where protection effectiveness has been recently evaluated: Belize (e.g., Acosta 2006; Chan et al. 2013; Acosta et al. 2019), the Bahamas Exuma Cays Land and Sea Park (e.g., Kough 2023; Kough et al. 2019), Florida Keys (e.g., Delgado and Glazer, 2020), and Serrana Bank (Colombia, e.g., Ardila et al. 2020).

### 2.5 Model evaluation

To validate our suitability predictions, we evaluated model estimations for the current scenario (2000-2020) and the 2020-2040 interval using iNaturalist occurrence data from 2000-2020 and 2020-2025, respectively (GBIF 2026). We used only ‘research grade’ observations and applied a distance-based thinning procedure, retaining a single record per environmental pixel to reduce spatial autocorrelation (similarly to the process described above). Additionally, to reduce uncertainty, observations falling outside the species’ reported depth range (McCarthy 2007) were excluded. We evaluated the predictive power of our models using a binomial significance test for ecological niche models, implemented in the ntbox R package (Osorio-Olvera et al. 2020). To minimize spatial non-independence between calibration and evaluation datasets, we restricted model testing to occurrences located in pixels not used during calibration. While this approach does not ensure absolute independence, it reduces the redundancy of environmental information and mitigates inflated performance estimates (see Pitfall 5 in Anderson 2012).

### 2.6 Model uncertainty

To evaluate uncertainty in our model projections in regard to the environmental divergence between calibration and transfer scenarios, we conducted a mobility-oriented parity (MOP) analysis using the functions available for this analysis in kuenm2. This method measures the multivariate environmental distance between the background points in the calibration area (reference) and each pixel within the transfer (future-climate) scenarios. We restricted this analysis to the environmental PCs used in the final models, and we further constrained it by the species’ known depth range. By quantifying similarity between reference and transfer conditions, we identified potential non-analogous climates where environmental variables, individual, multiple, or in combination, fall outside the range of calibration data. This allowed us to detect areas of strict extrapolation, where similarity values tend toward zero, indicating higher uncertainty in the projected suitability (Owens et al. 2013; Cobos et al. 2024).

## 3 Results

Of the 1170 candidate models tested using Maxent, one met the selection criteria. Similarly, only one of the 26 models tested was retained following model calibration and evaluation using ellipsoids (Supplementary Data 1 and Supplementary Tables 1-2). For Maxent models, the distribution and extent of suitable areas varied depending on the extrapolation scheme (see Figures 2-5 and Supplementary Figures 2-4). Suitability estimations from both frameworks (Maxent and ellipsoids) for the present period (2000-2020) produced similar distributions of suitable areas along the northern coast of South America, the insular Caribbean, and parts of Central America (Figure 2). However, notable differences occurred in the Gulf of Mexico and along the southern coast of the US, where ellipsoid models predicted narrower suitable areas than Maxent models. Approximately 92% of the total pixels containing benthic seagrass habitats in the Caribbean were identified as suitable by both Maxent and ellipsoid models (Supplementary Figure 5).

**Figure 2:**
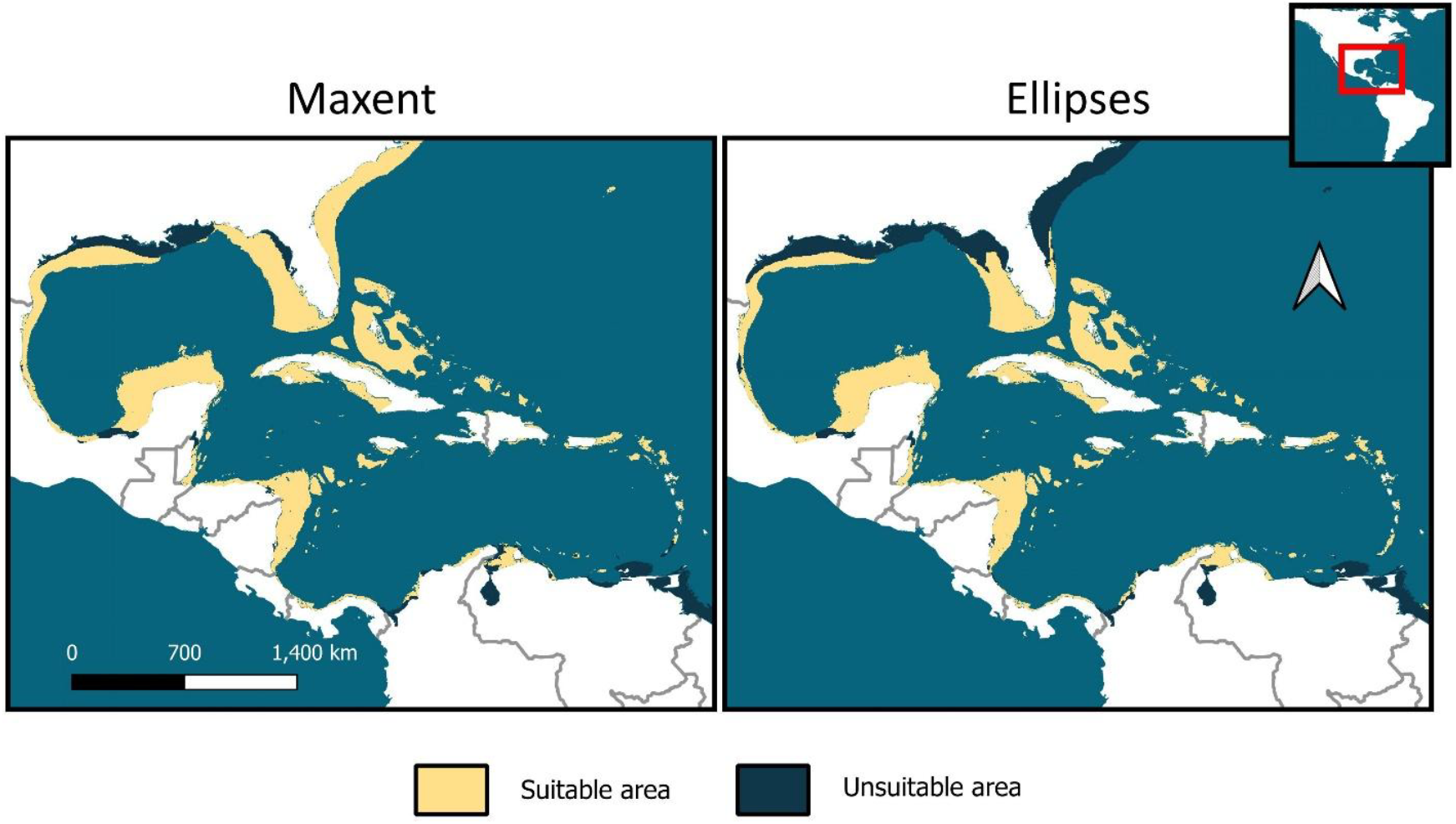
Estimated suitable areas for the queen conch (*Aliger gigas*) under the current scenario (2000-2020). Predictions produced using Maxent with free extrapolation (left) and ellipsoid models (right).

The rate of suitable area loss per 20-year interval varied across climate change scenarios. Under the most conservative scenario (SSP-126), both Maxent and ellipsoid models predicted similar proportions of area loss, although they differed in the specific regions where these losses were concentrated. Both modeling frameworks estimated consistent patterns of loss of currently suitable areas along the coasts of South America, Central America, and the Greater Antilles. The Maxent model estimated more pronounced losses than ellipsoid models in the west coast of Florida, Gulf of Mexico, Yucatan Peninsula and the Bahamas. Areas projected by both frameworks to retain suitability for the longest timeframe were concentrated along the southeastern coast of the United States, the northern coast of the Yucatan Peninsula, northern and northeastern Bahamas, parts of Turks and Caicos, and northwestern Gulf of Mexico (Figure 3).

**Figure 3:**
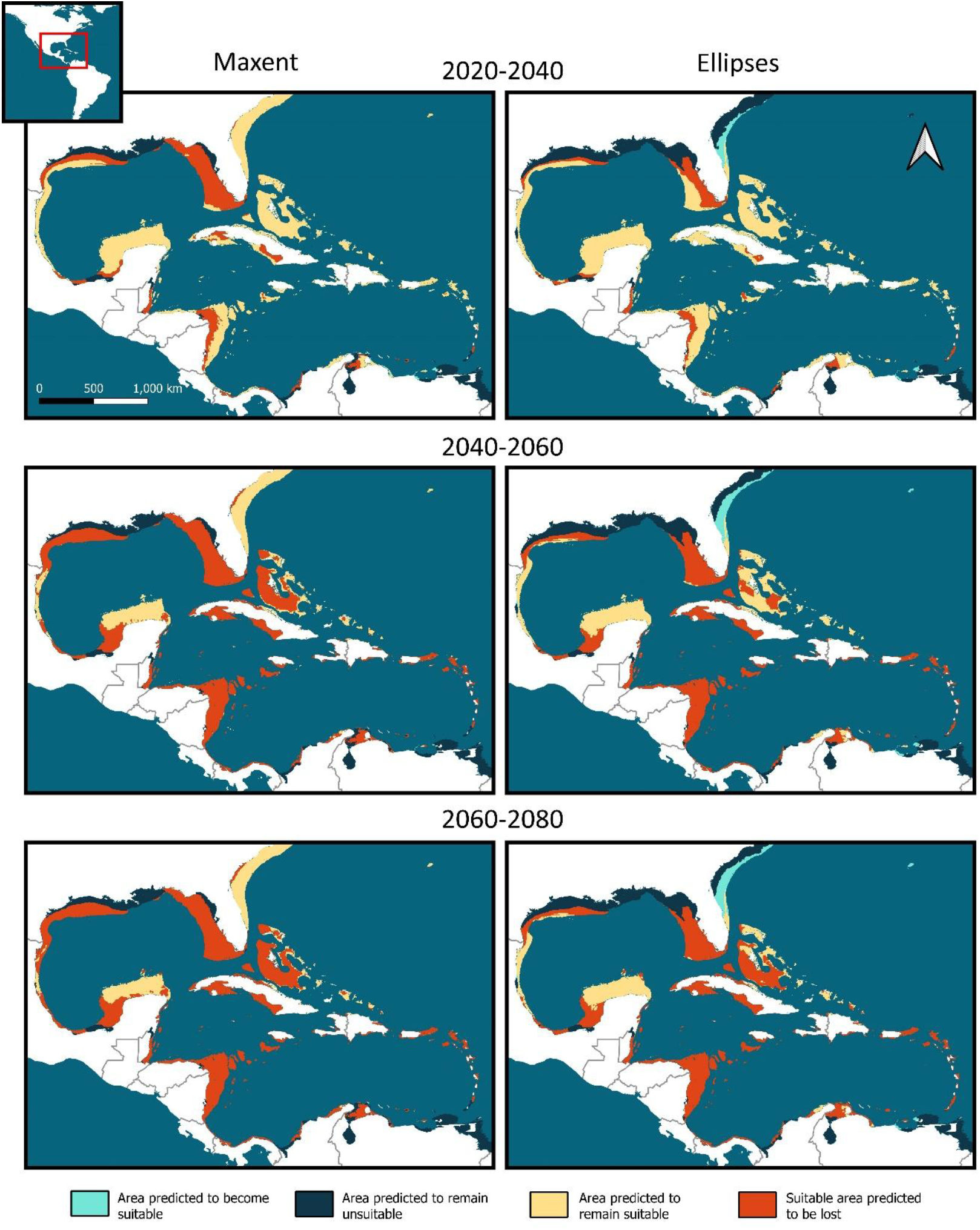
Projected changes in environmental suitability for the queen conch (*Aliger gigas*) under the Shared Socioeconomic Pathway 126 (SSP-126) scenario. Rows represent changes from the present day through three future time intervals. Suitability predictions were generated using Maxent with free extrapolation (left) and ellipsoid models (right).

Under the moderate climate change scenario (SSP-370), both Maxent and ellipsoid models projected a complete loss of abiotically suitable areas across the Caribbean by the 2040-2060 period. According to our models, areas currently suitable in the Gulf of Mexico, along the southeastern coast of the United States, throughout the Greater and Lesser Antilles and along fragmented portions of the Central America and South America coasts are projected to persist through 2020-2040 under both modeling frameworks. Both frameworks also projected that limited suitable areas along the coasts of Florida, Georgia, and South Carolina would persist or emerge during the 2040–2060 period. No suitable areas were projected beyond 2060 under this scenario (Figure 4).

**Figure 4:**
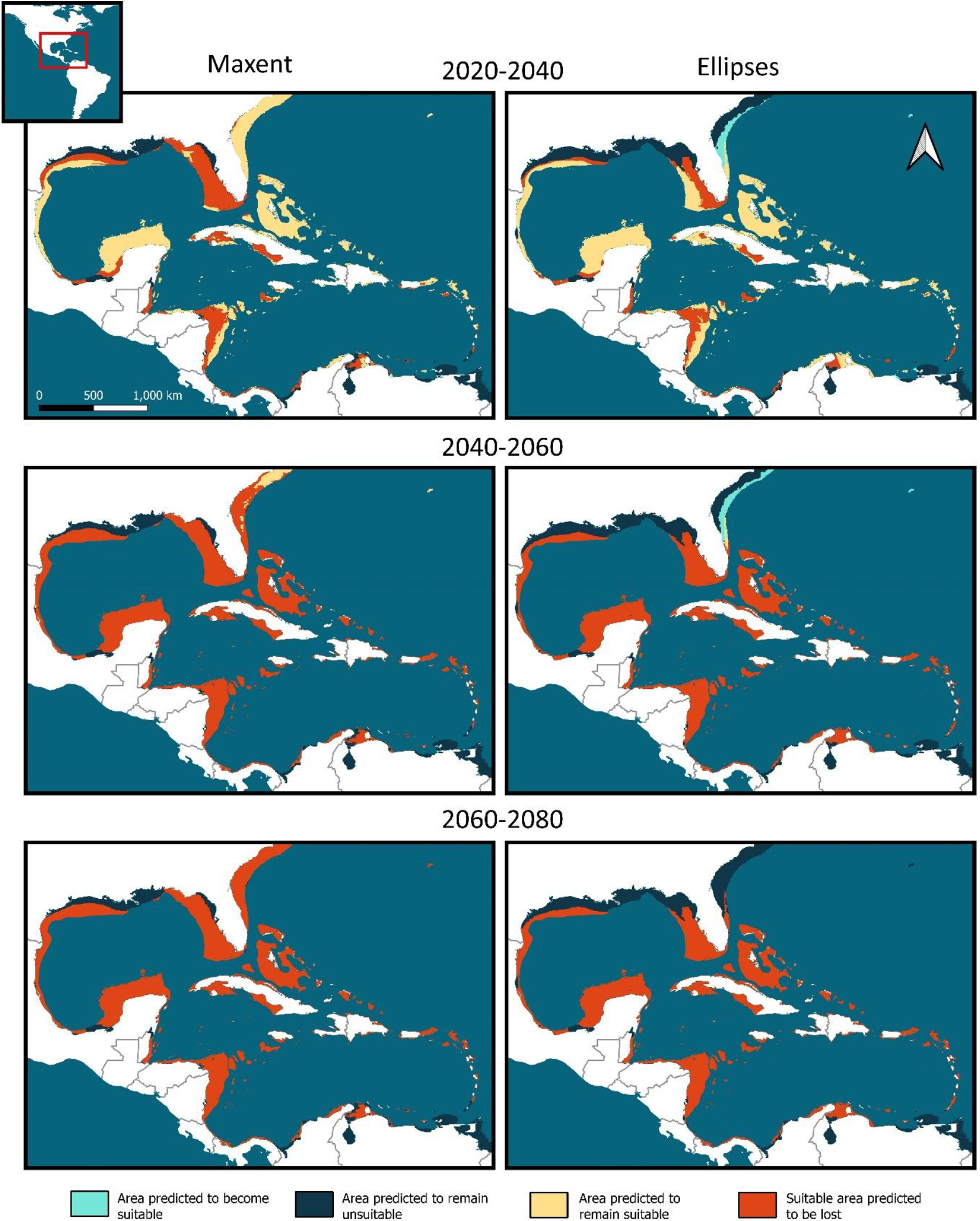
Projected changes in environmental suitability for the queen conch (*Aliger gigas*) under Shared Socioeconomic Pathway 370 (SSP-370) scenario. Rows represent changes from the present day through three future time intervals. Suitability predictions were generated using Maxent with free extrapolation (left) and ellipsoid models (right).

Both modelling frameworks exhibit parallel trends across the moderate and extreme (SSP-585) emission scenarios but with an accelerated rate of suitable area contraction (Figure 5). Under this scenario, suitable areas along the southeastern coast of the United States projected by both modeling frameworks in the moderate scenario were further reduced (Figure 5).

**Figure 5:**
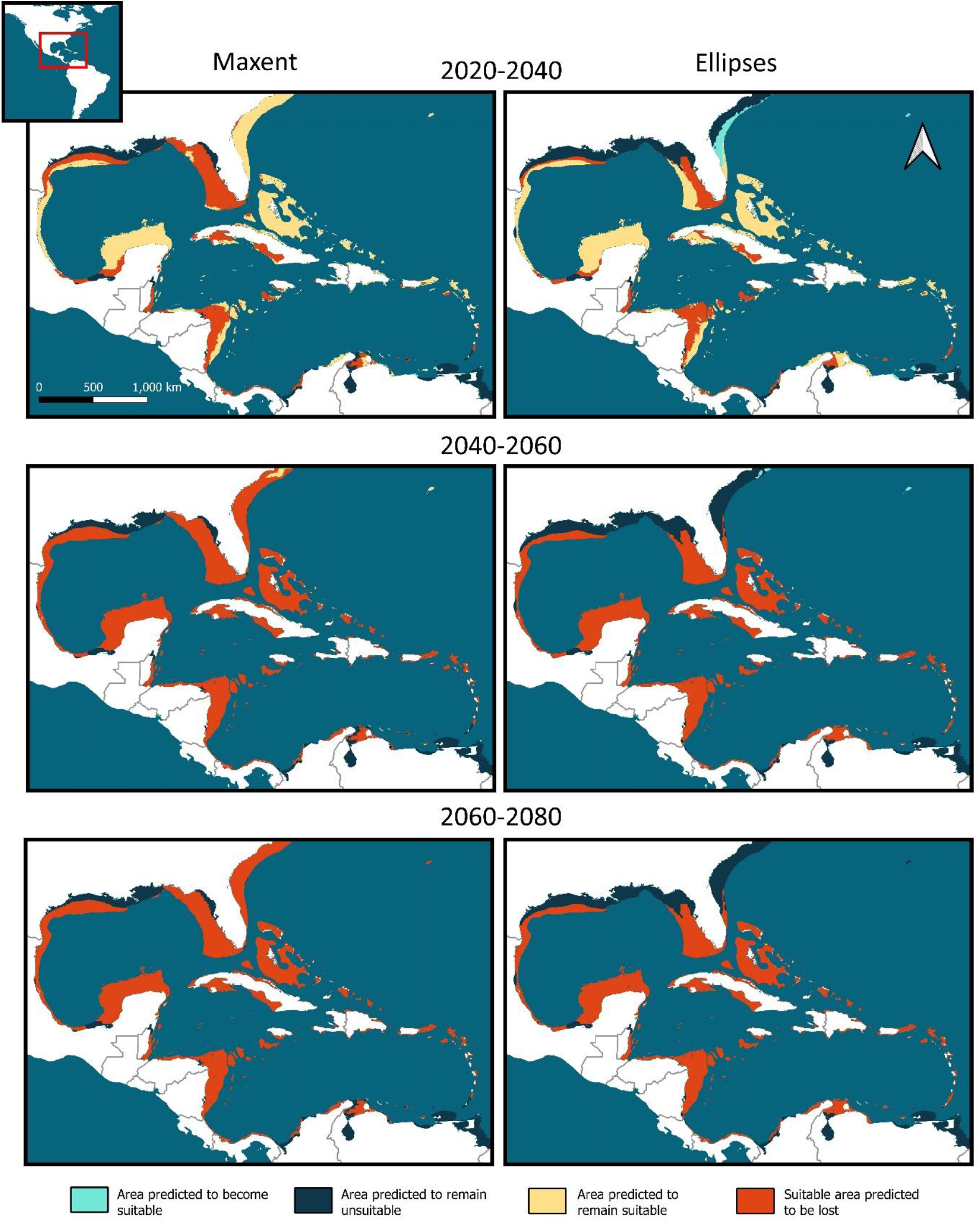
Projected changes in environmental suitability for the queen conch (*Aliger gigas*) under the Shared Socioeconomic Pathway 585 (SSP-585) scenario. Rows represent changes from the present day through three future time intervals. Suitability predictions were generated using Maxent with free extrapolation (left) and ellipsoid models (right).

All models yielded statistically significant estimates (Supplementary Tables 3-18), that is, they predicted presence more often than expected by chance relative to the proportion of total map pixels. When validated using iNaturalist data for the 2000–2010 period, both frameworks demonstrated high predictive power (true positive rates: Maxent ∼95%, ellipsoids ∼97%; see Figure 6). However, when validated against recent iNaturalist data (2020–2025), ellipsoid models showed equal or higher predictive power than Maxent models across all climate change scenarios (Supplementary Tables 5-10 and 13-18). Maxent models performed poorly primarily across Florida, Belize, Honduras, Panama, Colombia and Venezuela, as well as to a lesser extent in the Puerto Rico and southern Lesser Antilles. Conversely, ellipsoid models exhibited higher overall accuracy and lower commission rates, which were localized primarily to Florida, the Yucatán Peninsula, the Panamanian coast, and the southern Lesser Antilles (Figure 7).

**Figure 6:**
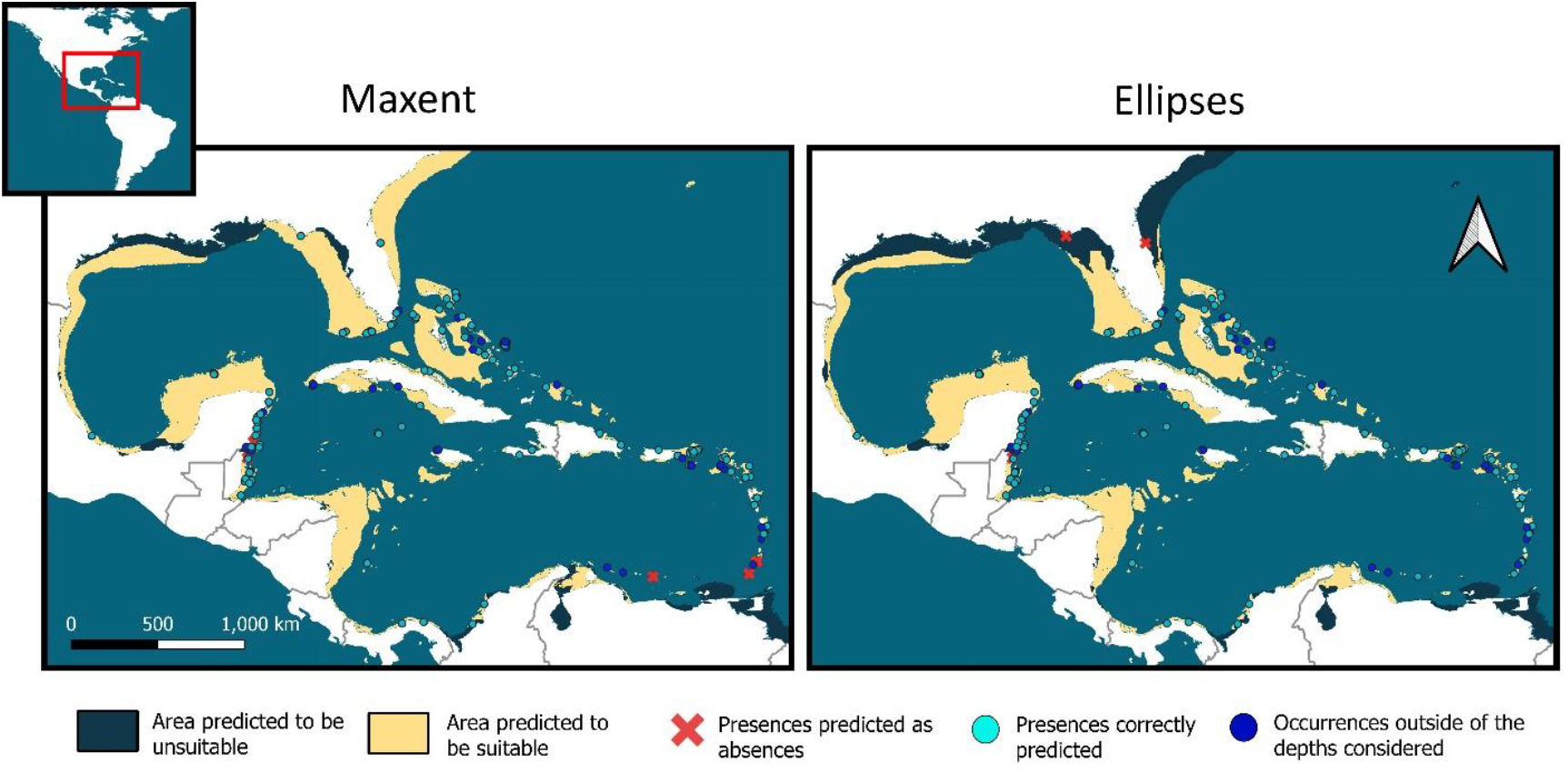
Performance evaluation of suitability predictions for the current scenario (2000-2020). Suitability predictions were generated using both Maxent with free extrapolation (left) and ellipsoid models (right). Evaluation was done using “Research grade” iNaturalist occurrence data from 2000-2020.

**Figure 7:**
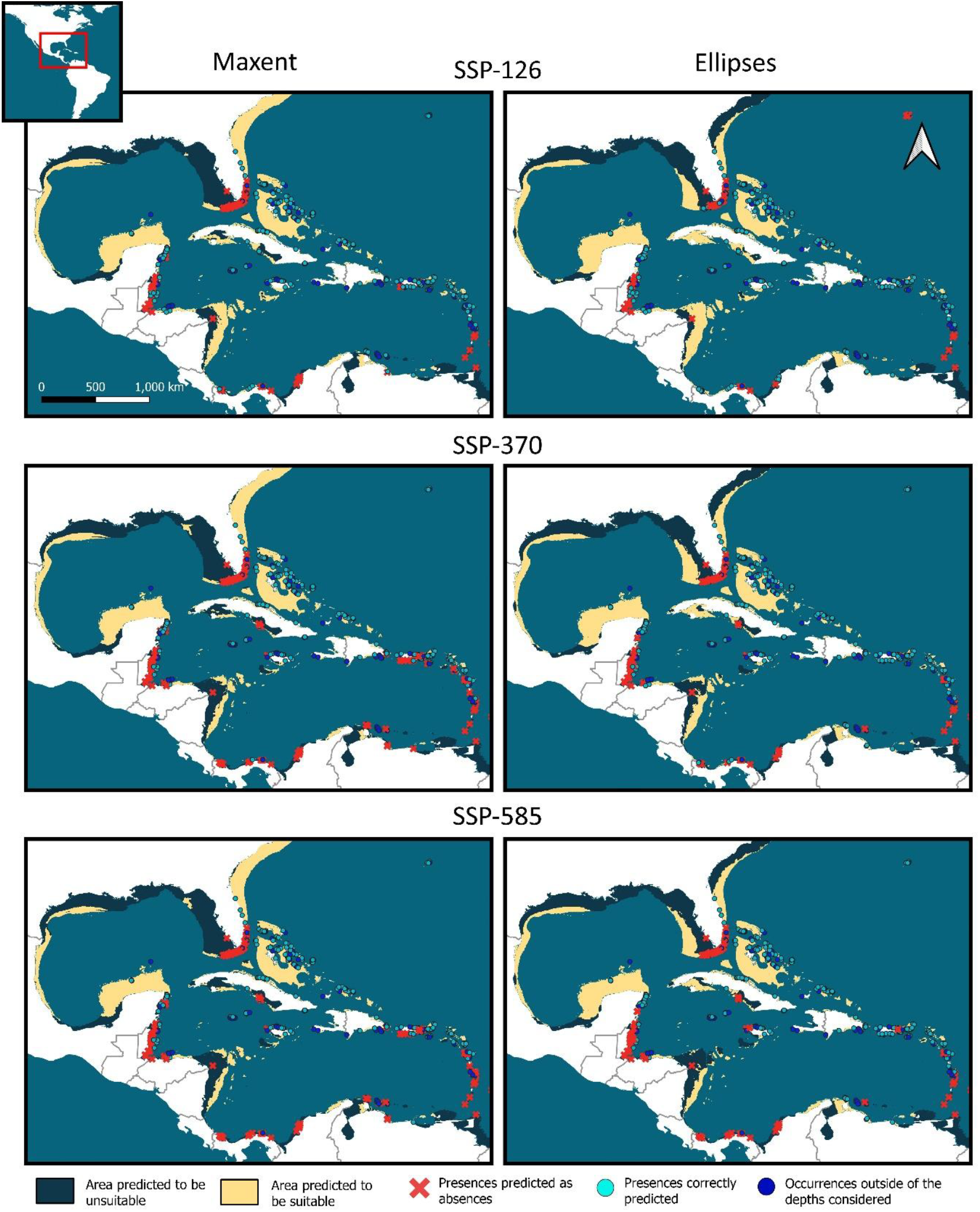
Performance evaluation of suitability predictions for the period 2020-2040 under the three Shared Socioeconomic Pathways scenarios considered. Predictions produced using both Maxent with free extrapolation models (left) and ellipses (right). Evaluation was done using “Research grade” iNaturalist occurrence data from 2000-2020.

As regards the current configuration of MPAs in the Caribbean region in relation to queen conch protection, our results suggest that only ∼15% of the total present-day suitable area is covered by protected areas (14% for Maxent and 16.70% for ellipsoids estimations, see Supplementary Data 2-3). Furthermore, even under the most optimistic scenario (SSP-126), only one of the four MPAs of special emphasis, the Exuma Cays Land and Sea Park, are projected to retain abiotically suitable areas through 2060-2080 (see Supplementary Figure 1C-D). Under the moderate and extreme scenarios (SSP-370), the Exuma Cays Land and Sea Park and part of the western edge of the Florida Keys Marine Sanctuary are expected to remain partially suitable until the 2020-2040 period. None of the four MPAs of special emphasis are projected to maintain abiotically suitable areas for the species beyond 2040 under these scenarios.

Assessments of strict extrapolation risk showed considerable differences between the calibration area and some of the future scenarios considered herein (Supplementary Figures 6-7). Extrapolation toward higher PC1 and PC2 values was observed in the central and southern Caribbean, as well as along portions of the Texas and Louisiana coasts. In contrast, extrapolation toward lower values was primarily associated with PC4, and to a lesser extent PC1, within the northern Caribbean. Overall, extrapolation and its associated uncertainty toward higher values increased southward in scenarios farther into the future, particularly under the more extreme scenarios. On the other hand, uncertainty toward lower values increased northward in more extreme scenarios that were increasingly distant in time.

## 4 Discussion

We modeled current and future potential suitable areas for queen conch across the Caribbean using ecological niche models. Our findings indicate a consistent northward shift in suitable areas over the coming decades, a trend that persists across all climate scenarios and modeling frameworks.

Several currently suitable areas in South America, Central America, and the Lesser Antilles are projected to become unsuitable. These projections coincide with regions where population declines have been reported in recent decades (e.g., Tewfik and Guzmán 2003; Acosta 2006; De Jesús-Navarrete and Valencia-Hernández 2013; Wynne et al. 2016; Marco et al. 2021; Osorto 2025; Ministerio de Ambiente y Desarrollo Sostenible [Minambiente] et al., 2025). These regions are also areas where some of the lowest adult densities (i.e., individuals per unit area) have been documented (Horn et al. 2022; Vaz et al 2022). Although the drivers of present and future population declines may differ—ranging from environmental factors (e.g., temperature, salinity) and anthropogenic pressures (e.g., overfishing of larval source populations, habitat destruction) to demographic constraints (e.g., Allee effect)—populations likely become more vulnerable as their environments become less suitable. More generally, the fact that modeled suitability loss is anticipated in regions already experiencing population declines raises significant concerns about the long-term survival of the queen conch in these areas.

Our results reiterate the importance of seagrass beds for queen conch populations. Although these results are consistent with the current understanding of the species’ habitat requirements—as seagrass provides both nutrition and protection from predators during juvenile stages (Ray and Stoner 1995; Stoner 2003; Stoner et al. 2011)—our estimates are not without limitations. The association between juvenile conch and seagrass cover is not exclusive; many seagrass meadows, particularly turtlegrass (*Thalassia testudinum*), have been reported to be unoccupied by juveniles (Stoner 2003). This suggests that additional environmental, oceanographic, or biotic factors influence habitat suitability within these beds, requiring local-scale evaluations to identify these factors. Furthermore, the spatial estimates of seagrass used in this study do not represent *T. testudinum* exclusively; therefore, the actual distributional range of this species may be smaller than the seagrass extent estimated herein. Consequently, our models may overestimate suitability, particularly given recent findings of higher growth probabilities for juvenile conch in native compared to invasive seagrass species (Boman et al. 2019). Finally, adult queen conchs occupy a broader range of habitats beyond seagrass (Stoner 2003; Stoner and Davis 2010), including sandy bottoms and coral rubble (McCarthy 2007). Because this study relies primarily on adult presence data and was conducted at an environmental resolution of ∼5 km, our estimates may not accurately capture habitat use during earlier life stages or at finer spatial scales (e.g., see case of Exuma Sound in The Bahamas described in Stoner 2003 and Stoner and Ray 1993, 1996).

Our findings also provide deeper insights into queen conch distribution when analyzed in the context of population connectivity and larval transport dynamics, particularly regarding the location, size, and isolation of suitable habitat patches. For this species, larval exchange between populations is largely mediated by surface currents in the Caribbean (Stoner 2003; Vaz et al. 2022; Horn et al. 2022). Thus, the presence of suitable habitats along these current pathways is critical for facilitating larval settlement, population establishment, and gene flow among existing populations.

Our models identified several areas projected to remain suitable until at least 2040 under the optimistic scenario (SSP-126). Among these, the northern Caribbean, including the northern coast of the Yucatán Peninsula, regions of the Gulf of Mexico, and parts of Florida, the Bahamas, and Turks and Caicos, appears to be the most resilient in terms of abiotic suitability, while southern Caribbean regions appear to be the most vulnerable under the scenarios evaluated. Under the moderate and extreme scenarios, the contraction of suitable areas accelerates markedly over time, with areas of abiotic suitability expected to persist only through 2020–2040. Connectivity modeling system (CMS) simulations developed by Vaz et al. (2022) indicate that southeastern Caribbean populations are likely disconnected from those in the western and northwestern Caribbean, primarily due to localized overfishing in critical corridors such as Puerto Rico, the Dominican Republic, and Haiti. The recent discovery of a deep-water nursery habitat in Puerto Rico (Cruz-Marrero et al. 2024) could have important implications for regional connectivity, though its contribution to restoring it remains to be assessed. This disjunction nonetheless suggests that northern populations, such as those in Florida, the Bahamas and Turks and Caicos, currently rely almost exclusively on self-recruitment rather than larval transport from Central America (Nicaragua, Belize, and Mexico) and parts of the Greater Antilles (Dominican Republic and Haiti; Vaz et al. 2022). Consequently, some areas identified here as abiotically suitable in the northwestern Caribbean and Central America may host either self-recruiting populations (see Delgado et al. 2008 for specific evidence of this in Florida) or, if densities are too low for sustaining successful reproduction (see Figure 6 in Horn et al. 2022, p. 27), non-reproducing populations, as appears to be the case in Belize, Mexico, and Florida (see Figure 7 in Vaz et al. 2022). Recent evidence from the Florida Keys further corroborates the influence of density-dependent reproduction on population persistence (Delgado and Glazer, 2020). Thus, while some of these regions may hold the longest-persisting areas of abiotic suitability over time, factors such as local population densities and larval transport dynamics will likely determine the long-term persistence of populations.

Suitable areas predicted across different warming scenarios and modeling frameworks—despite varying rates of loss—include the southeastern coast of the United States, the western coast of Florida, and the Gulf of Mexico (Figures 2-5). However, the probability of populations establishing or persisting in these areas is likely constrained by oceanographic factors mediating larval transport (Stoner 2003; Vaz et al. 2022). Although abiotically favorable, these regions face low recruitment potential because prevailing ocean currents typically transport surface waters through the Strait of Florida toward the northeastern Atlantic (see Figure 1 in Vaz et al. 2022 for a detailed visualization of sea currents in the Caribbean). A similar dynamic occurs in the northwestern Gulf of Mexico, where conditions are predicted to be suitable, but currents flowing northward may reduce the likelihood of larval settlement. These patterns underscore that within abiotically favorable areas, long-term persistence is strongly influenced by oceanographic dynamics. Interestingly, queen conch has been reported in some of these regions historically (Supplementary Figure 8), suggesting that natural populations may still exist or could reestablish despite reduced larval input relative to other Caribbean areas. Consequently, further evaluation of these suitable areas may be valuable to assess their potential for hosting translocated populations, especially given the success of previous translocation efforts (e.g., Delgado et al. 2004).

Currently, only 14-17% of present-day abiotically suitable areas for queen conch are covered by MPAs in the Caribbean, and our models project a substantial contraction of these areas over the next 40 years. These results underscore the urgent need for new protected areas that prioritize abiotically suitable regions where populations remain at optimal sizes and densities (e.g., Bahamas, Cuba, Turks and Caicos, and Colombia, see abundance estimates in Horn et al. 2022 and Ardila et al. 2020) and serve as critical larval sources (see larval sources in Vaz et al. 2022). Notably, part of the Exuma Cays Land and Sea Park (Bahamas) is expected to remain suitable until 2060-2080 under the optimistic climate change scenario considered herein, while southern MPAs in Serrana Bank (Colombia) and Belize may become abiotically unsuitable as early as 2040-2060. This finding highlights the uncertainty regarding the long-term capacity of northern MPAs to sustain queen conch populations without larval input from southern sources. Restoring connectivity and potentially establishing new populations through seeding from southern MPAs while they remain abiotically suitable, will be critical for ensuring long-term conservation of the species.

Although our analyses are quantitatively robust, our results are not without limitations. First, the performance of Maxent models for the 2020-2040 period was notably poor in Florida, Central America, South America, and parts of the Lesser Antilles. Therefore, these specific regional predictions should be interpreted with caution. In contrast, ellipsoid models demonstrated greater reliability through higher predictive power and closer alignment with recent occurrence records, yet they did not entirely overcome the predictive challenges identified in the aforementioned regions. Second, strict model extrapolation and associated uncertainty in certain areas of the Caribbean increased under more extreme climate scenarios and in scenarios farther into the future. Accordingly, suitability estimates for the 2040–2060 and 2060–2080 periods—particularly under moderate and extreme climate scenarios—should be interpreted in light of these limitations. Third, our results are based on 20-year environmental statistics, which may limit the model’s ability to detect suitability patterns driven by finer-scale temporal climatic variability. Fourth, even though we consider one of the most important and well-known biotic interactions of juvenile queen conch with species of seagrasses, we did not incorporate other relevant interactions, such as predation and competition, that could influence distribution at other life stages (e.g., larval stage). Finally, our analyses did not consider potential anthropogenic factors that may impact the distribution, such as fishing zones, marine pollution, coastal development and tourism. Incorporating these variables into future spatial analyses would further refine distribution estimates and help prioritize sites for population conservation and translocation.

This study provides the first regional-scale estimate of queen conch (*Aliger gigas*) distribution using an ecological niche modeling framework. Rather than serving as another attempt to map the species’ known range, our work builds upon and enhances existing knowledge about the long-term survival of queen conch populations throughout the Caribbean. By integrating our models with current understanding of larval transport, population connectivity, reproductive biology, and life-history, we offer a more comprehensive perspective on potential distributional shifts over the coming decades. Ultimately, this research highlights the value of ecological niche modeling as a critical, complementary tool for informing biodiversity conservation strategies and assessing the risk to the long-term survival of the queen conch.

## 5 Data Availability Statement

The data and code scripts used/generated in this study can be found in the OSF repository: https://osf.io/gvhda/overview?view_only=f16586131b944f8ca9cc8e04b0cd8028.

## 7 Author Contributions

DRA: Conceived and designed the study, developed the methodology, performed the analyses, and contributed to the interpretation and discussion of results and the design of figures. CNP: Contributed to the development of methodology, data analysis, interpretation and discussion of results, and the design of figures. ZRU: Performed the MOP analysis and contributed to the development of methodology, discussion of results, and the design of figures. All authors contributed to manuscript review and approved the submitted version.

## 9 Acknowledgments

We thank the members of the KUENM group at the University of Kansas, whose discussions and feedback greatly benefited this study. We especially thank Marlon Cobos and Townsend Peterson for their thorough and constructive feedback on the manuscript and the theoretical and methodological framework of the study.

---

**Figure S1.**
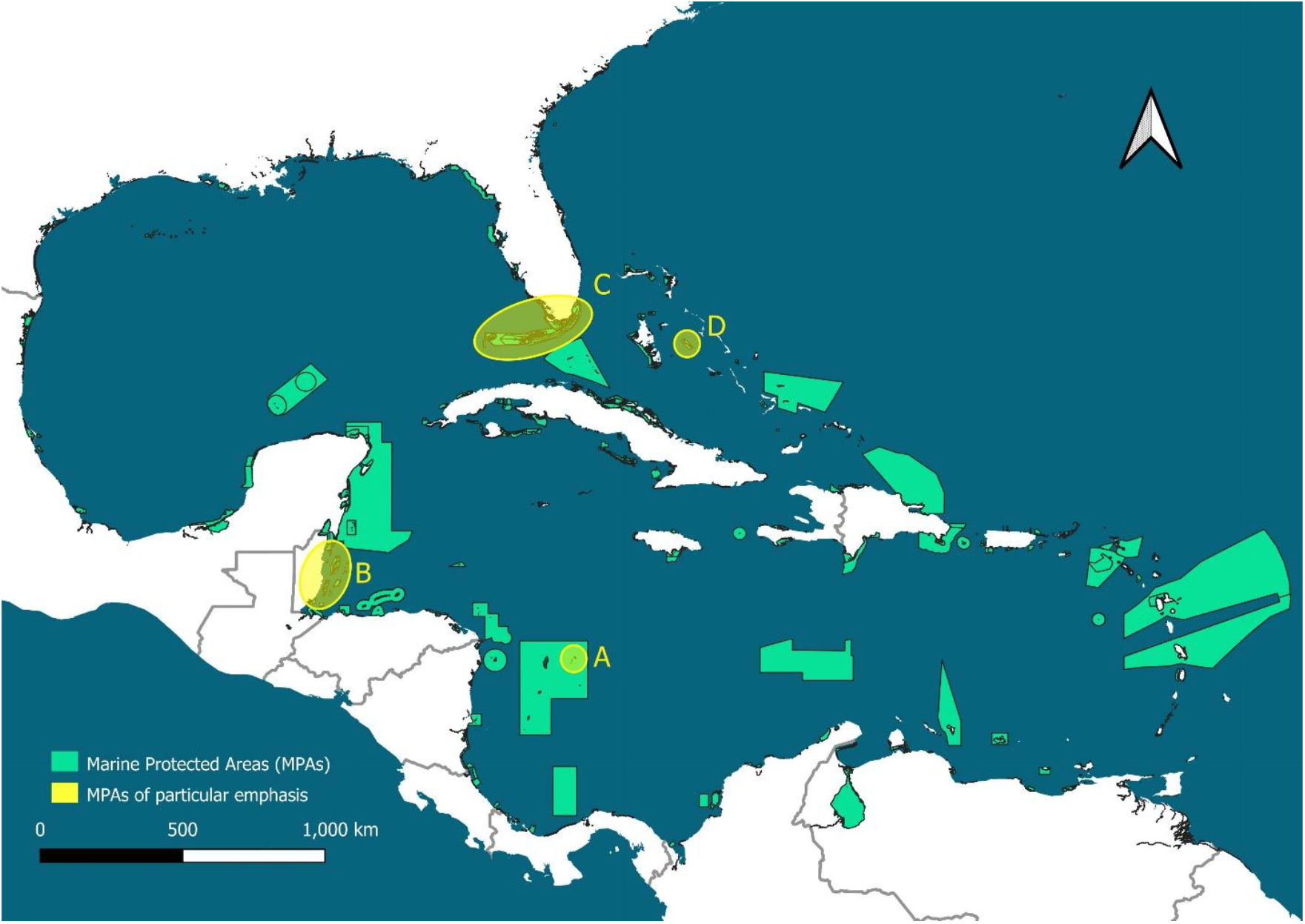
Marine protected areas (MPAs) in the Caribbean (light blue). Obtained from UNEP-WCMC and IUCN (2024). Four MPAs of particular emphasis in this study are indicated by yellow circles. A) Serrana Bank (Seaflower MPA, Colombia), B) Belize MPAs, C) Florida Keys Marine Sanctuary, D) Exuma Cays Land and Sea Park.

**Figure S2.**
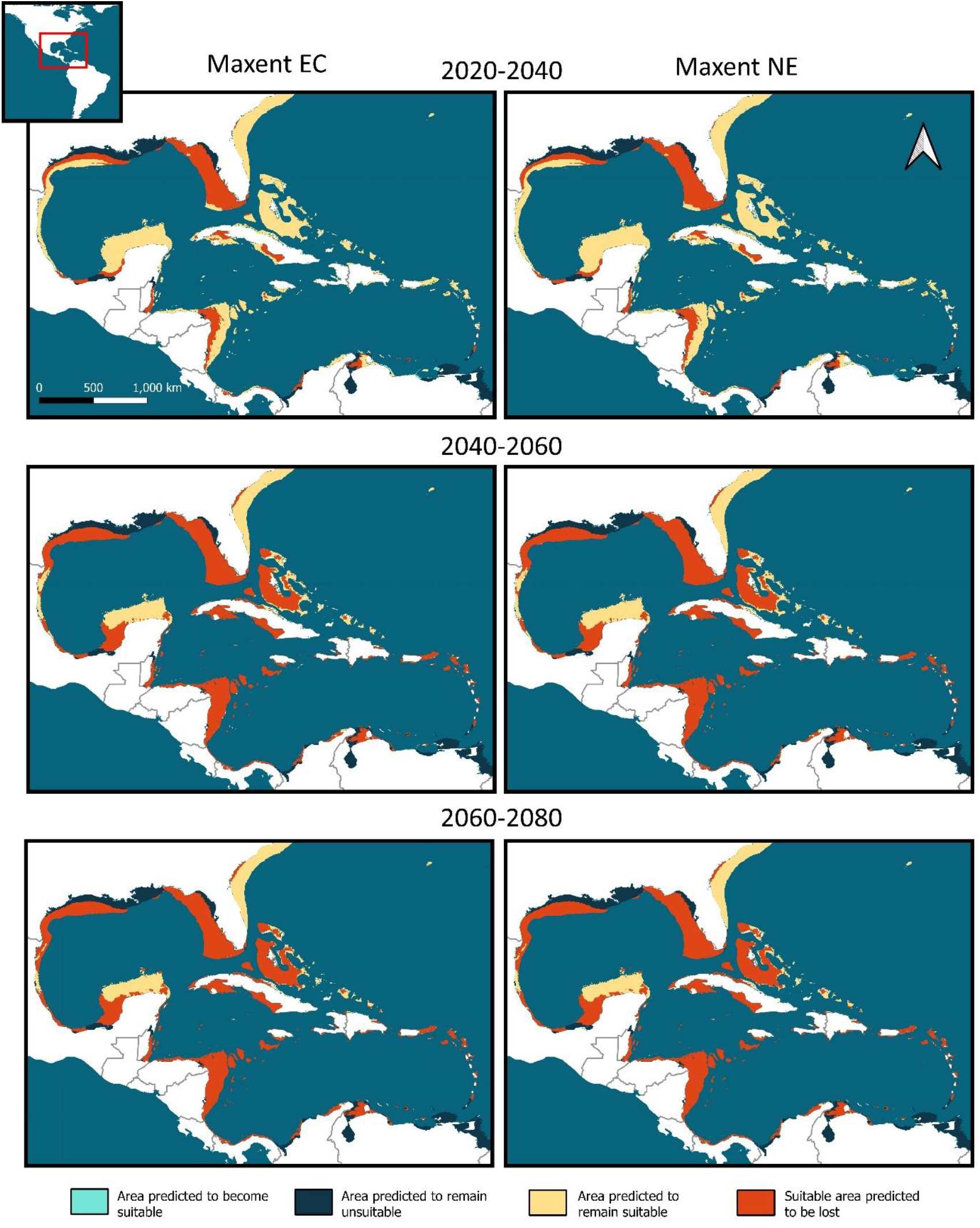
Marine protected areas (MPAs) in the Caribbean (light blue). Obtained from UNEP-WCMC and IUCN (2024). Four MPAs of particular emphasis in this study are indicated by yellow circles. A) Serrana Bank (Seaflower MPA, Colombia), B) Belize MPAs, C) Florida Keys Marine Sanctuary, D) Exuma Cays Land and Sea Park.

**Figure S3.**
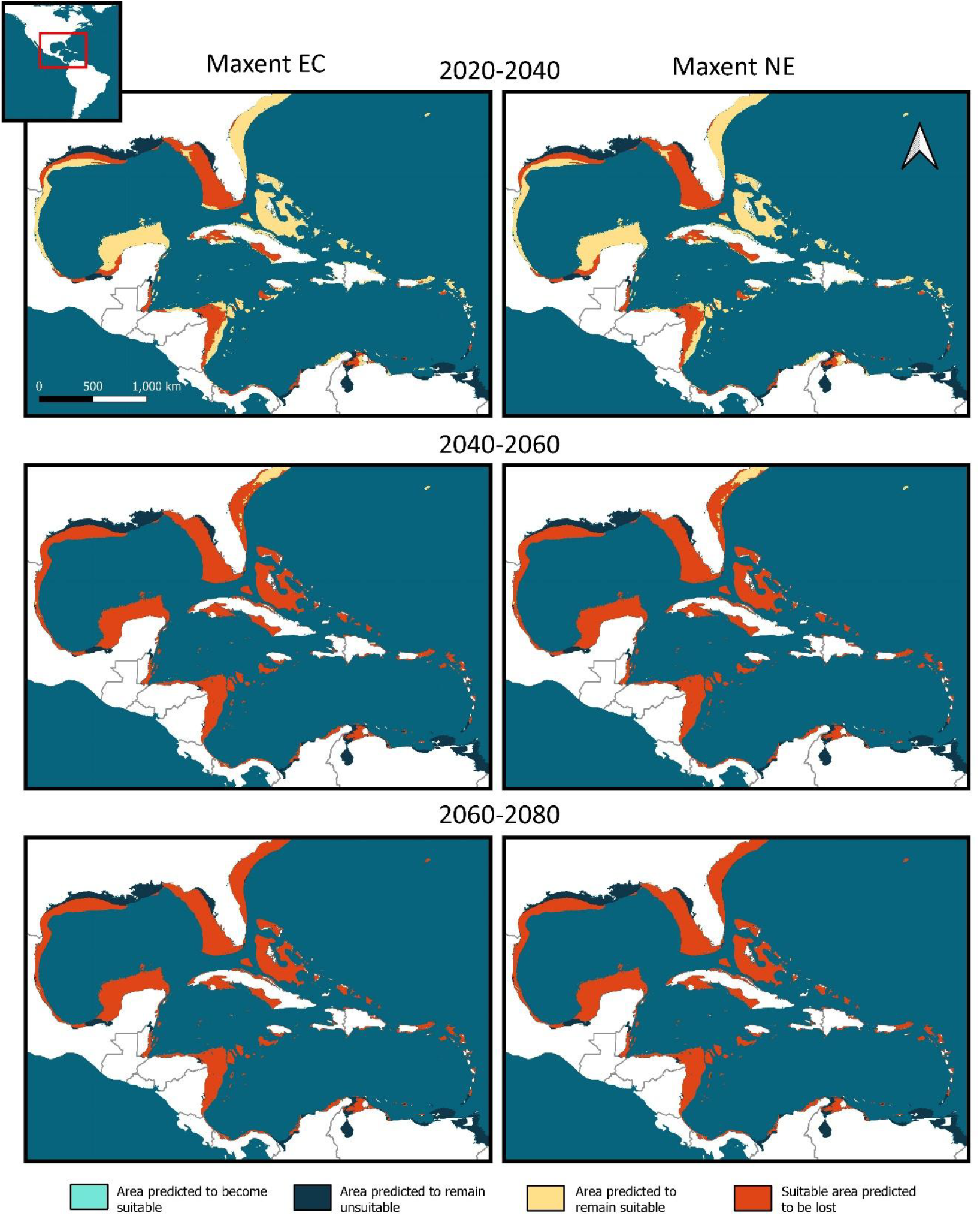
Projected changes in environmental suitability for the queen conch (*Aliger gigas*) under the Shared Socioeconomic Pathway 370 (SSP-370) scenario. Rows represent changes from the present day through three future time intervals. Suitability predictions were generated using Maxent models considering extrapolation with clamping (EC, left) and no extrapolation (NE, right).

**Figure S4.**
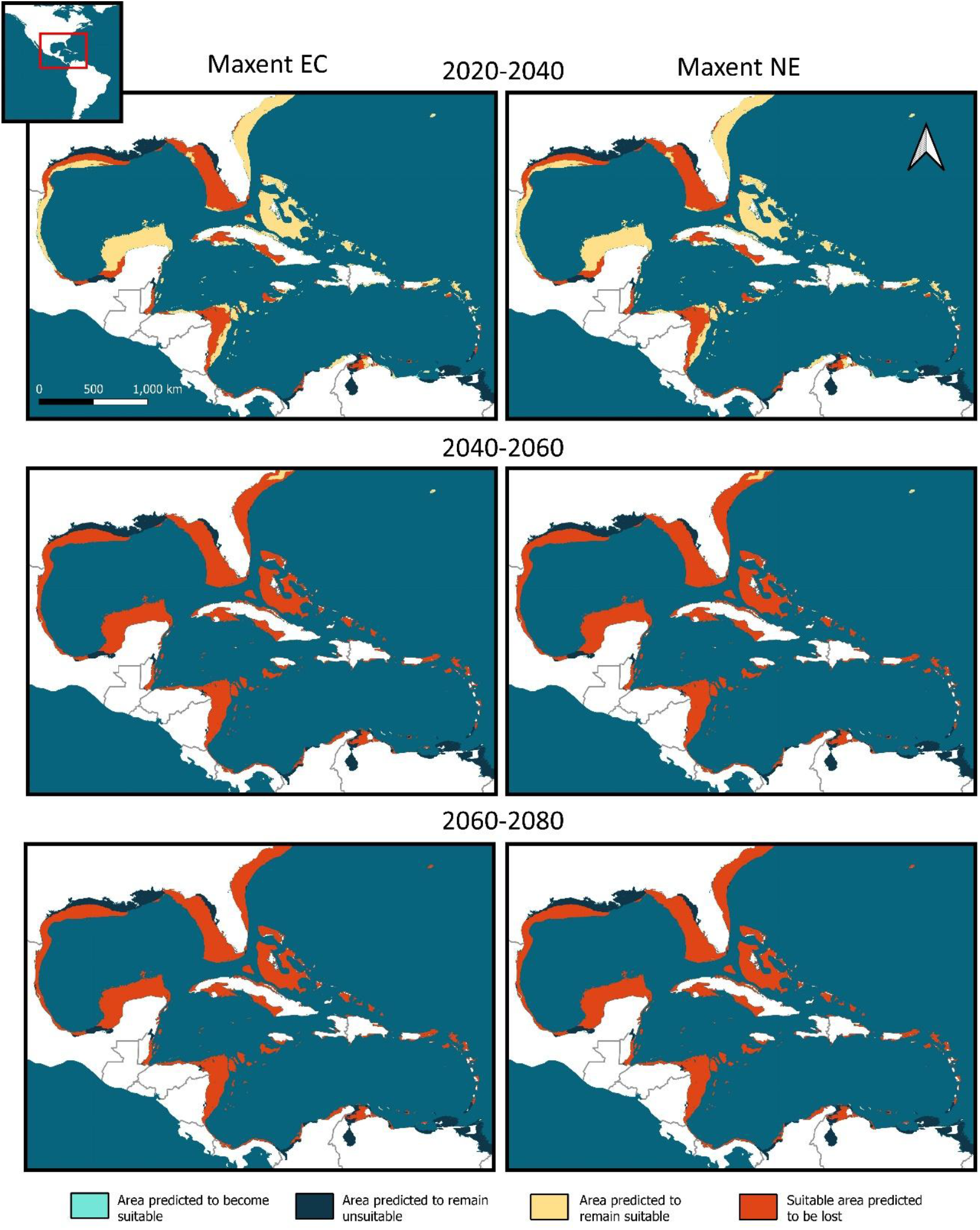
Projected changes in environmental suitability for the queen conch (*Aliger gigas*) under the Shared Socioeconomic Pathway 585 (SSP-585) scenario. Rows represent changes from the present day through three future time intervals. Suitability predictions were generated using Maxent models considering extrapolation with clamping (EC, left) and no extrapolation (NE, right).

**Figure S5.**
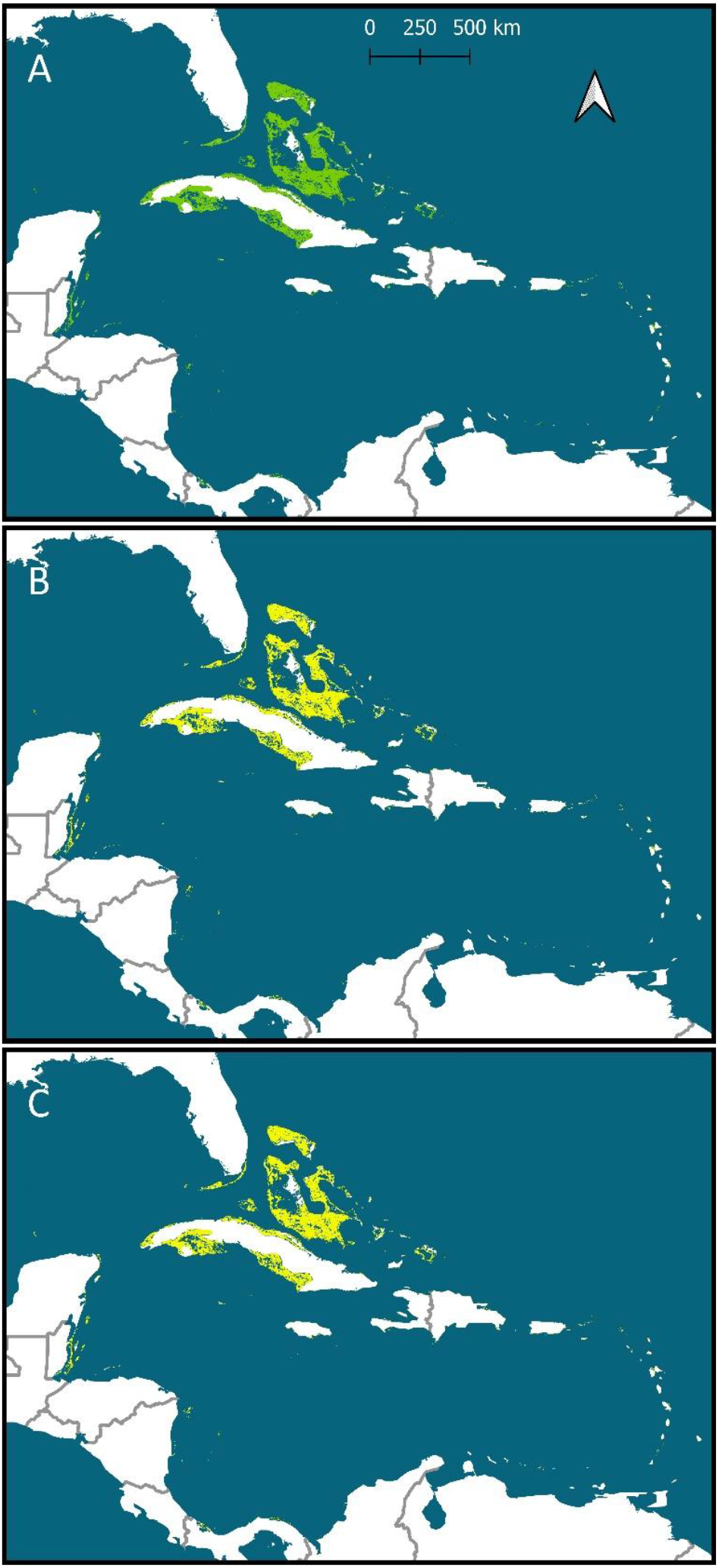
A) Total coverage of benthic seagrass habitats in the insular Caribbean (Schill et al. 2021) highlighted in green. B) Total coverage of benthic seagrass (green) and currently suitable seagrass habitat predicted by our Maxent model (yellow). C) Total coverage of benthic seagrass (green) and currently suitable seagrass habitat predicted by our ellipsoid model (yellow).

**Figure S6.**
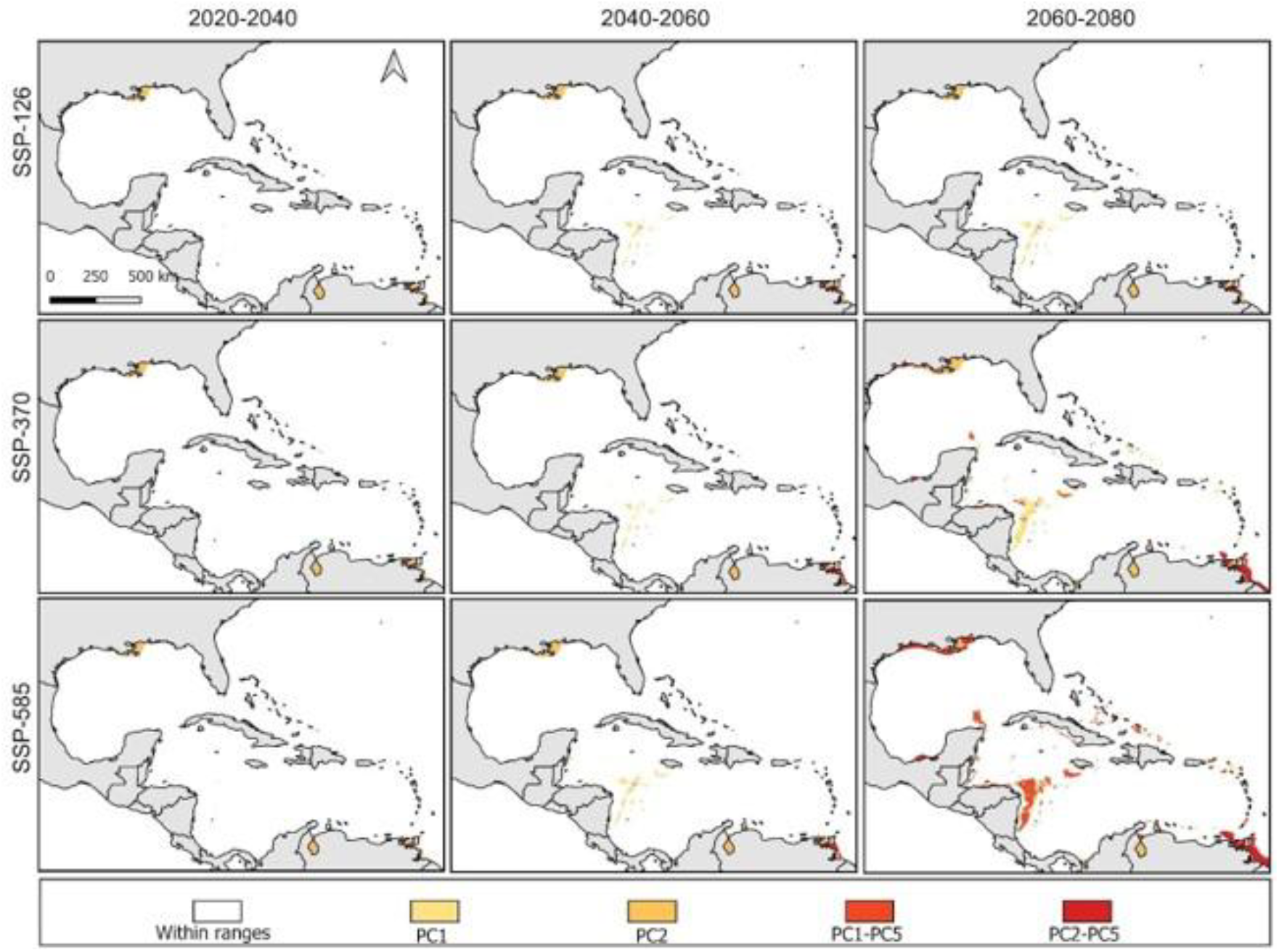
Areas exhibiting a risk of strict extrapolation toward higher environmental values during model transfer due to non-analogous conditions in specific variables.

**Figure S7.**
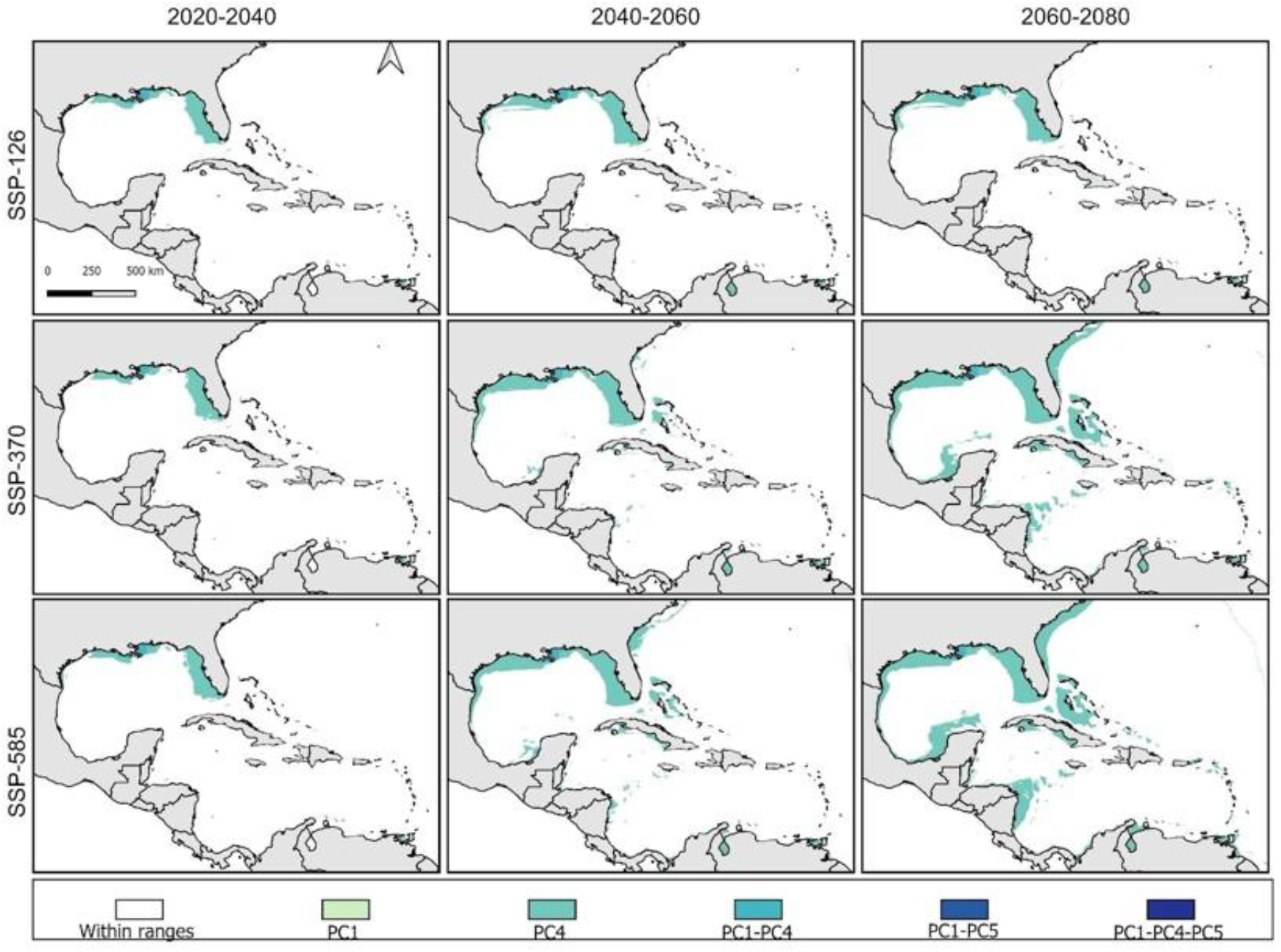
Areas exhibiting a risk of strict extrapolation toward lower environmental values during model transfer due to non-analogous conditions in specific variables.

**Figure S8.**
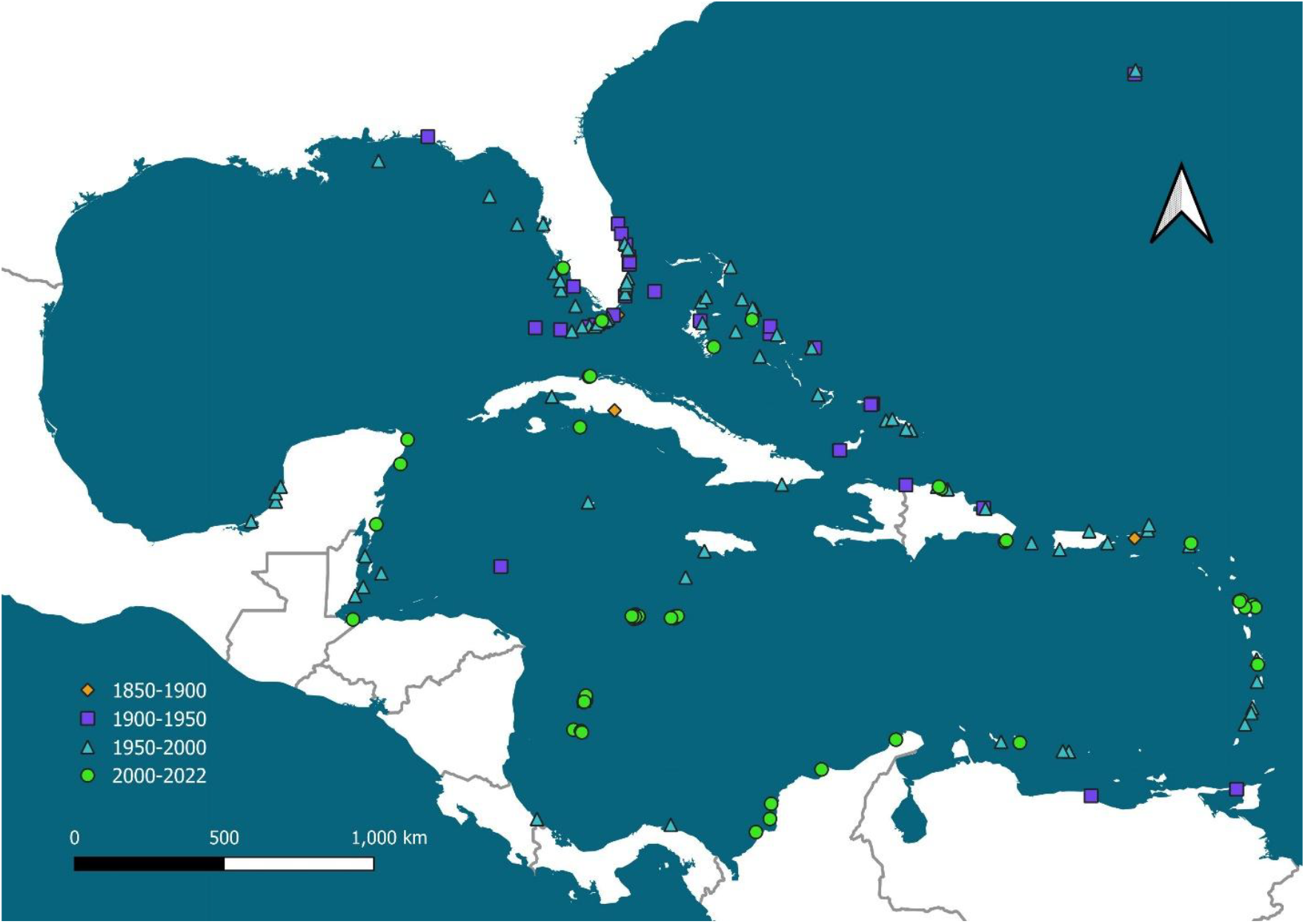
Clean occurrence records for queen conch between 1850 and 2020 in our dataset.

